# The transmission-blocking effects antimalarial drugs revisited: mosquito fitness costs and sporontocidal effects of artesunate and sulfadoxine-pyrimethamine

**DOI:** 10.1101/2020.05.19.103408

**Authors:** M Villa, M Buysse, A Berthomieu, A Rivero

## Abstract

Assays used to evaluate the transmission-blocking activity of antimalarial drugs are largely focused on their potential to inhibit or reduce the infectivity of gametocytes, the blood stages of the parasite that are responsible for the onward transmission to the mosquito vector. For this purpose, the drug is administered concomitantly with the gametocyte-infected blood, and the results are evaluated as the % reduction in the number of oocysts in the mosquito midgut.

We report the results of a series of experiments that explore the transmission blocking potential of two key antimalarial drugs, artesunate (AS) and sulfadoxine-pyrimethamine (SP), when administered to mosquitoes already infected from a previous blood meal. For this purpose, uninfected mosquitoes and mosquitoes carrying a 6-day old *Plasmodium relictum* infection (early oocyst stages) are allowed to feed either on a drug-treated or an untreated host in a fully factorial experiment. This protocol allows us to bypass the gametocyte stages and establish whether the drugs are able to arrest the ongoing development of oocysts and sporozoites, as would be the case when a mosquito takes a post-infection treated blood meal. In a separate experiment, we also explore whether a drug-treated blood meal impacts key life history traits of the mosquito relevant for transmission, and if this depends on their infection status.

Our results show that feeding on an AS- or SP-treated host has no epidemiologically relevant effects on the fitness of infected or uninfected mosquitoes. In contrast, when infected mosquitoes feed on an SP-treated host, we observe both a significant increase in the number of oocysts in the midgut, and a drastic decrease in both sporozoite prevalence (−30%) and burden (−80%) compared to the untreated controls. We discuss the potential mechanisms underlying these seemingly contradictory results and contend that, provided the results are translatable to human malaria, the potential epidemiological and evolutionary consequences of the current preventive use of SP in malaria-endemic countries could be substantial.

## 1. Introduction

Synthetic antimalarial drugs are the mainstay for the prevention and treatment of malaria throughout the world. Over the last century, these synthetic antimalarials, have replaced traditional herbal remedies such as the (quinine-containing) bark of the South American cinchona trees and the (artemisin-containing) Chinese drug *qinghao*. Broadly speaking, four major synthetic antimalarial drug classes exist for the treatment of malaria: (i) quinolines (chloroquine, mefloquine), which owe their origins to quinine, interfere with the ability of *Plasmodium* detoxify hematin, a toxic compound resulting from the degradation of the haemoglobin; (ii) antifolates (pyrimethamine, sulfadoxine, proguanil), so-called because they disturb the folate pathway of *Plasmodium*, thereby interfering with its DNA and amino-acid synthesis; (iii) atovaquone which interferes with the parasite’s mitochondrial electron transport, and (iv) artemisinin derivatives (artemether, artesunate), the most potent and effective anti-malarials to date, which exert their anti-malarial action by perturbing redox homeostasis and the haematin detoxification in the parasite (Müller and Hyde, 2010).

Although the prime purpose for developing these antimalarials is obviously to prevent or cure the infection of the patients, it has become rapidly obvious that they can also be used to reduce the prevalence of the disease in the population by reducing the onward transmission of the parasite by the vector (Sinden et al., 2012; Wadi et al., 2019). The transmission-blocking effect of antimalarial drugs can take place in three different, albeit non-exclusive, ways. Firstly, drugs may be able to kill, arrest the maturation, alter the sex ratio or reduce the infectivity of gametocytes, the sexual stages of the parasite that are present in the blood and are responsible for the transmission to the mosquito. Secondly, drugs may be able to hinder the development of the parasite within the mosquito. *Plasmodium* development inside the mosquito is complex and involves the fusion of male and female gametocytes to form a *zygote*, the passage of the mobile zygote through the midgut wall to form an *oocyst* that grows, undergoing successive mitosis, ruptures and releases thousands of *sporozoites* that migrate to the salivary glands. Antimalarial drugs, or their metabolites, can find their way to the mosquito midgut where they can block the parasite either directly, by being toxic to any of the above stages of the parasite, or indirectly, by disturbing the fine-tuned mosquito physiological pathways that are essential for parasite development (Sinden et al., 2012). Finally, just as some antimalarials have unwanted secondary effects in the host, so they may be able to adversely affect key life history traits of the mosquito essential for parasite transmission such as its longevity or host seeking behaviour.

To date, the majority of experimental studies have focused on the gametocytocidal effects of antimalarial drugs (Delves et al., 2018; Ruecker et al., 2014). Gametocytes are an attractive target for transmission-blocking interventions because they constitute an important bottleneck in the life cycle of the parasite and can be directly targeted due to their presence in the bloodstream of the host. While some of the available compounds may be able to achieve a 100% gametocyte inhibition, and thus completely block the transmission cycle, many of the compounds being tested result in a partial, if at times substantial, gametocyte reductions (Sanders et al., 2014). Partial reductions may, however, fail to accurately reflect the degree to which the mosquitoes become infected (Churcher et al., 2013; Sinden, 2017), and may even end up enhancing transmission due higher mosquito survival rates associated to lower oocyst burdens (Sinden, 2010). It has therefore been argued that interventions that aim to target gametocytes should be combined with others that target the later stages of parasite development within the mosquito (Blagborough et al., 2013; Paaijmans and Fernàndez-Busquets, 2014; Sinden, 2010). Whether antimalarial drugs have an effect on oocyst or sporozoite development, however, is still largely unknown, as most protocols provide the drug with the infected blood meal thereby conflating the effects on the gametocytes with the effects on the later stages (Delves et al., 2018; Wadi et al., 2018).

Current WHO advice for the treatment of malaria in endemic countries relies heavily on the use of two synthetic antimalarial drugs: artesunate and sulfadoxine-pyrimethamine (WHO, 2019). Artesunate (henceforth AS), a potent and fast-acting artemisinin derivative, is used as a treatment for severe/complicated malaria, or in combination with longer-acting antimalarial drugs such as sulfadoxine-pyrimethamine, amodiaquine or mefloquine for the treatment of children and adults with uncomplicated malaria (WHO, 2019). AS is a pro-drug that, once inside the cell, is rapidly converted into its active form, dihydroartemisinin, which in turn generates reactive oxygen species (ROS) that increase oxidative stress and cause malarial protein damage via alkylation (Hou and Huang, 2016; O’Neill et al., 2010). In humans, peak plasma concentrations are reached in 1–2 h following oral intake (though the absorption through injection may be slower and somewhat more variable, Balint, 2001). Sulfadoxine-pyrimethamine (henceforth SP), on the other hand, is recommended in areas with moderate to high malaria transmission for the intermittent preventive treatment (ITP) of pregnant women and infants (0-1 years) and, in areas of high seasonal transmission, for the seasonal malaria chemoprevention (SMC) of young children (<6 years of age, WHO, 2019). Each year, millions of people throughout the world get treated by one of these drugs (WHO, 2019). Sulfadoxine and pyrimethamine act synergistically to inhibit the activity of dihydropteroate synthase (DHPS) and dihydrofolate reductase (DHFR), respectively, thus inhibiting the folic acid metabolism of the parasite (Peterson et al., 1988). They are both long-lasting drugs, with plasma concentrations being found up to 42 days post treatment (Karunajeewa et al., 2009).

The gametocytocidal potential of both drugs has been the subject of numerous studies (review in Butcher, 1997; Wadi et al., 2019). AS has a demonstrated cytocidal activity against mature and immature gametocytes *in vitro* (Chotivanich et al., 2006; Peatey et al., 2012). Evidence of the gametocytocidal effects of SP, on the other hand, is contradictory. Certain studies have found that SP has no gametocidal activity (Miguel-Blanco et al., 2015; Plouffe et al., 2016), others that SP inhibits male gametocyte formation (Delves et al., 2012, 2013) and yet others that pyrimethamine administration results in an increased gametocyte production, possibly as an adaptive response of the parasite to stressful conditions (Buckling et al., 1999).

Here, we report the results from a series of experiments that aim to investigate the transmission-blocking potential of AS and SP downstream from their putative cytocidal effects on the gametocytes. For this purpose, we perform a series of factorial experiments feeding infected and uninfected mosquitoes either on drug-treated or control hosts. Experiments are carried out using the avian malaria parasite, *Plasmodium relictum* and its natural vector, the mosquito *Culex quinquefasciatus*, one of the few available systems in which these experiments are both technically and ethically possible. In order to bypass the gametocytocidal effects of the drug, the ‘infected’ mosquitoes were exposed to the drug while carrying a 6-day old infection from a previous blood meal (corresponding the early oocyst stages, Pigeault, 2015). Under our standard laboratory conditions, 5-6 days is the average length of the *Cx quinquefasciatus* gonotrophic cycle.

Avian malaria has played a key historical role in the study of human malaria, being a stimulus for the development of medical parasitology (Rivero and Gandon, 2018). It has played a particularly pivotal role in the screening and clinical testing of the first synthetic antimalarials (Coatney et al., 1953; Hewitt, 1940; Rivero and Gandon, 2018) and in the study of their potential use as transmission-blocking compounds (Gerberg, 1971; Ramakrishnan et al., 1963; Terzakis, 1971). Compared with rodent malaria, the avian malaria system has the added advantage of using the parasite’s natural vector in the wild, the mosquito *Culex pipiens*, thereby sidestepping the issues associated with mosquito-parasite combinations without a common evolutionary history (Cohuet et al., 2006; Dong et al., 2006).

Our aims were to establish: 1) whether the drugs administered to infected mosquitoes can arrest the ongoing development of oocysts and/or sporozoites, but also 2) whether the drugs alter the fitness of mosquitoes and, if so, whether this is contingent on whether the mosquitoes are infected with *Plasmodium*. Our results provide insights into the multiplicity of effects that a given drug may have in the different stages of the parasite’s sporogonic cycle. We discuss the potential epidemiological and evolutionary consequences of using AS and SP to reduce transmission of *Plasmodium* in the field.

## 2. Material and Methods

### 2.1. Mosquito and parasite protocols

All experiments were carried out using a laboratory strain of *Culex pipiens quinquefasciatus* (SLAB strain). *Culex* mosquitoes are the most important natural vector of avian malaria in Europe and the Americas. The larvae in all the experiments were reared at a constant density per tray (n=300 larvae) following previously published laboratory protocols (Vézilier et al., 2010). Larval trays (n=22) were placed individually inside an “emergence cage” (40 cm x 28 cm x 31 cm) and emerged adults were allowed to feed *ad libitum* on a 10% glucose water solution. Rearing and experiments took place at our standard insectary conditions (24-26 °C, 60-80% RH, and 12:12 L:D photoperiod).

*Plasmodium relictum* (lineage SGS1) is the aetiological agent of the most prevalent form of avian malaria in Europe. The parasite lineage was isolated from blue tits (*Parus caeruleous*) collected in the Montpellier area in October 2016 and subsequently passaged to naïve canaries (*Serinus canaria*) by intraperitoneal injection. Since then, it has been maintained by carrying out regular passages between our stock canaries through intraperitoneal injections with the occasional passage through the mosquito.

### 2.2. Impact of antimalarials on Plasmodium-infected and uninfected mosquito traits

The purpose of these experiments was to establish whether feeding from a sulfadoxine-pyrimethamine (SP) or an artesunate (AS) treated host can negatively influence mosquito traits such as longevity and fecundity. For this purpose, two separate experiments were set up.

#### 2.2.1. Sulfadoxine-pyrimethamine experiment

To obtain infected and uninfected mosquitoes to use in the experiment, 200 female mosquitoes were placed in a cage containing either an infected or an uninfected bird (n=4 and n=3 cages of infected and uninfected birds respectively). Infected birds were obtained by injecting them with 100µL of blood from our *P. relictum-*infected canary stock. Mosquito blood feeding took place 10 days after the injection, to coincide with the acute phase of the *Plasmodium* infection in the blood (Cornet et al., 2014; Pigeault et al., 2015). After the blood meal, which took place overnight, the bird was taken out of the cage, unfed mosquitoes were discarded and engorged mosquitoes were provided with a 10% sugar solution. Three days later, a tray with water was placed inside the cage to allow egg laying (and hence the completion of the mosquito’s gonotrophic cycle). Seven days pbm 20 mosquitoes were haphazardly chosen from each of the 4 cages having contained an infected bird, and were dissected under a binocular microscope to verify the existence of *Plasmodium* oocysts in their midgut. These dissections confirmed that the large majority of the mosquitoes (91 %) had become infected.

To explore the impact of SP on the fecundity and longevity of mosquitoes, infected and uninfected mosquitoes were allowed to take a second blood meal on either an SP-treated or a control bird. For this purpose, four days prior to the blood meal, 3 birds (henceforth SP-treated birds) had a daily subcutaneous injection of 30 µl of a sulfadoxine-pyrimethamine solution (Sigma S7821 and 46706, 320 mg/kg Sulfadoxine, 16 mg/kg Pyrimethamine solubilized in DMSO) while 3 additional (control) birds were injected with 30 µl of DMSO. The red blood cell count of birds (number of red blood cells per ml of blood) was quantified immediately before the blood meal using flow cytometry (Beckman Coulter Counter, Series Z1). One hour after the last injection, 100 infected and 80 uninfected mosquitoes were placed in a cage containing either an SP-treated or a control bird. To allow the identification of the infected and uninfected mosquitoes, they were previously marked using a small amount (2.5 µg/female) of coloured fluorescent powder (RadGlo^®^ JST) as a dust storm. Preliminary trials have shown that, at this concentration, the dust has no effect on mosquito traits (Vézilier et al., 2012). On day 1 post blood meal (pbm), the number of blood-fed mosquitoes in each of the cages was counted and unfed females discarded.

To quantify haematin (a proxy for blood meal size) and fecundity, 80 females from each cage (40 infected and 40 uninfected) were haphazardly chosen and placed individually in numbered 30 ml Drosophila tubes, covered with a mesh (‘haematin tubes’). Food was provided in the form of a paper strip soaked in a 10% glucose solution. Three days later (day 4 pbm), all mosquitoes were transferred to a new tube containing 7 mL of mineral water to allow the females to lay their eggs (‘fecundity tube’). The amount of haematin excreted at the bottom of each tube was quantified as an estimate of the blood meal size following previously published protocols (Vézilier et al. 2010). The fecundity tubes were provided with a paper strip soaked with 10% sugar solution. The fecundity tubes were checked daily for the presence of eggs. The egg laying date was recorded and egg rafts were photographed using a binocular microscope equipped with a numeric camera. Eggs counted using the Mesurim Pro freeware (Academie d’Amiens, France).

To quantify longevity, the rest of the infected and uninfected mosquitoes were kept in the cages and provided with a tray of water for egg laying for the first 6 days. Survival of these mosquitoes was assessed daily by counting dead individuals lying at the bottom of each cage until all females died.

#### 2.2.2. Artesunate experiment

The protocol used was identical to the one used in the SP experiment with only a few minor modifications. Here, four days prior to the blood meal, 3 birds (henceforth AS-treated birds) had a subcutaneous injection of 50 µl of an artesunate solution (16 mg/kg artesunate, Sigma A3731, in a 50mg/kg bicarbonate solution) twice daily (9am and 6pm) while 3 additional (control) birds were injected with 50 µl of the bicarbonate solution. As in the previous experiment, mosquito dissections confirmed that the large majority of the *Plasmodium* mosquitoes (95 %) were indeed infected.

### 2.3. Impact of antimalarials on Plasmodium infection within the mosquito

The purpose of these experiments was to establish whether antimalarial drugs can have an effect on the development of *Plasmodium* within the mosquito. For this purpose, we allowed previously-infected mosquitoes to feed on either SP-treated, AS-treated or control birds (n=3 birds each). At the time of feeding, mosquitoes had been infected for 6 days from a previous blood meal. Protocols used to infect mosquitoes and treat the birds were identical to those used in the two previous experiments.

To assess the impact of the drugs in the blood meal on the *Plasmodium* parasites developing within the mosquitoes, 15-20 mosquitoes were haphazardly chosen from each cage at three different intervals: 8-9 days, 11-12 days and 14 days post-infection (corresponding to 2-3 days, 5-6 days and 8 days after the treated blood meal). Based on previous results (Pigeault, 2015) these intervals correspond to the expected peak oocyst numbers, start of sporozoite production and peak sporozoite production, respectively. At each of these time points, each mosquito was dissected to count the number of oocysts in the midgut under the microscope (as in Vézilier et al. 2010), and its head-thorax was preserved at -20°C for the quantification of the sporozoites. Sporozoites were quantified using real-time quantitative PCR as the ratio of the parasite’s *cytb* gene relative to the mosquito’s *ace-2* gene (Zélé et al. 2014). As in the other experiments a large majority of the mosquitoes were infected (82%-87%).

### 2.4. Statistical analyses

Analyses were carried out using the R statistical package (v3.4.4). The different statistical models used are described in the Supplementary Materials (Tables S1 & S2). The general procedure to build models was as follows: treatment (AS, SP, control), and infection status (infected/uninfected) were fitted as fixed explanatory variables. Birds were fitted as a random effect. Where appropriate, haematin and dissection day were introduced into the model as an additional fixed variable. Since we observed differences between the different plates used for the colorimetric quantification of the haematin (Vézilier et al., 2010) the models were fitted with the haematin residuals of a model containing haematin as a response variable and plate as a fixed explanatory variable. Maximal models, including all higher order interactions, were simplified by sequentially eliminating non-significant terms and interactions to establish a minimal model. The significance of the explanatory variables was establish using a likelihood ratio test (LRT) which is approximately distributed as a chi-square distribution (Bolker, 2008) and using p = 0.05 as a cut-off p-value.

Survival data were analyzed using Cox proportional hazards mixed effect models (coxme). Proportion data (blood-fed females, egg laying females, oocyst and sporozoite burden) were analyzed using mixed linear models and a binomial distribution. Response variables that were highly overdispersed (number of eggs per raft, oocyst burden) were analyzed using mixed negative binomial models (glmmTMB). *A posteriori* contrasts were carried out by aggregating factor levels together and by testing the fit of the simplified model using a LRT (Crawley, 2007). Because of the small number of replications, differences in red blood cell counts between the birds in the different treatments were tested using Kruskal-Wallis non-parametric tests.

### 2.5. Ethics statement

Bird manipulations were carried out in strict accordance with the “National Charter on the Ethics of Animal Experimentation” of the French Government. Experiments were approved by the Ethical Committee for Animal Experimentation established by the authors’ institution (CNRS) under the auspices of the French Ministry of Education and Research (permit number CEEA-LR-1051).

The authors declare no conflict of interests

## 3. Results

### 3.1. Impact of antimalarials on Plasmodium-infected and uninfected mosquito traits

#### 3.1.1. Sulfadoxine-Pyrimethamine (SP) experiment

The vast majority of mosquitoes (97-100%) blood fed, independently of whether they were provided with an SP-treated or a control bird (model 1, χ^2^= 0.1306, p = 0.7178) and of their infection status (model #, χ^2^=3.1086, p = 0.0779). The amount of blood ingested (quantified as the amount of haematine excreted) was also similar across experimental conditions (model 2, *treatment*: χ^2^ = 0.6545, p = 0.4185; *infection*: χ^2^ = 0.0001, p = 0.9802). There was no difference in the haematocrit of SP-treated and untreated birds (model 3, χ^2^ =0.4286, p = 0.5127).

The probability of laying an egg raft was overall very high (85-95%) except for infected mosquitoes feeding on control birds (65%, model 4, *treatment*infection*: χ^2^ = 11.372, p = 0.001). Overall, females having fed in SP treated birds laid eggs earlier than those fed on control birds (model 5, LR.stat = 10.243, p = 0.001). Egg laying date also depended on the interaction between blood meal size and the infection status of the mosquito (model 5, LR.stat = 7.2853, p = 0.007). While for infected females blood meal size had no impact on oviposition day, uninfected females who took larger blood meals laid eggs earlier than those who took a smaller blood meals (LR.stat = 6.7296, p = 0.0094). Mosquito fecundity (number of eggs per raft) decreased with egg laying date (model 6, χ^2^ = 15.808, p <0.001) but was independent of both treatment (model 6, χ^2^ = 0.2478, p = 0.6186) and infection status (model 6, χ^2^ = 0.1048, p = 0.7462),

Uninfected mosquitoes lived significantly longer than their infected counterparts (model 7, HR ± se = 0.8167 ± 0.0766; χ^2^ = 7.4768, p = 0.006). This effect was however independent on whether the host had been previously treated with SP or not (model 7, χ^2^ = 0.0557, p = 0.8134). The results were identical when analyzing survival to day 14, the time at which sporozoite production peaks (model 8).

#### 3.1.2. Artesunate (AS) experiment

As above, the vast majority of mosquitoes (95-98%) blood fed, independently of whether they were provided with an AS-treated or a control bird (model 9, χ^2^ = 2.4543, p = 0.1172) and of their infection status (model 9, χ^2^ = 0.1085, p = 0.7418). The amount of blood ingested was also similar across experimental conditions (model 10, *treatment*: χ^2^ =0.0009, p = 0.4185; *infection*: χ^2^ = 0.787, p = 0.375). There was no difference in the haematocrit of AS-treated and untreated birds (model 10, χ^2^ = 1.4727, p = 0.2888).

The probability of laying an egg raft was overall very high (81-88%). As in the SP experiment, infected mosquitoes had a slightly lower chance of laying eggs than their infected counterparts, though here this effect was independent of whether they had fed on a treated or an untreated bird (model 11, χ^2^ = 4.3911, p = 0.0361). For mosquitoes feeding on AS-treated birds, the probability of laying an egg raft depended heavily on the amount of blood ingested: treated females that took a small blood meal saw their probability of laying eggs significantly reduced (mean ± s.e probability of egg laying for treated females in the lowest blood meal quartile: 55.4 ± 6.7 %, in the highest blood meal quartile: 92.6 ± 3.0 %). No such difference was found in mosquitoes that fed on untreated birds (lowest quartile: 78.2 ± 5.6 %, highest quartile 85.5 + 4.8 % ; model 11, *treatment*haematine*: χ^2^ = 8.0323, p = 0.0046, see Supplementary Materials, Figure S1). The egg laying date was independent of the treatment (model 11, LR.stat = 0.203, p= 0.6523) but was negatively correlated with the size of the blood meal: females that take smaller blood meals laid eggs later (model 12, LR.stat = 12.498, p < 0.001). Mosquito fecundity (number of eggs per raft) increased with blood meal size (model 13, χ^2^= 36.875, p < 0.001) but was independent of both treatment (model 13, χ^2^ = 0.2784, p = 0.5978) and infection status (χ^2^ = 0.9796, p = 0.3223).

Neither the artesunate treatment (model 14, χ^2^ = 0.0577, p = 0.8102) nor the mosquito infection status (model 14, χ^2^ = 0.3266, p-value = 0.5677) had an impact on overall mosquito survival. The results were identical when analyzing survival to day 14, the time at which sporozoite production peaks (model 15).

### 3.2. Impact of antimalarials on Plasmodium infection within the mosquito

#### 3.2.1. Sulfadoxine-Pyrimethamine experiment

The prevalence of oocysts decreased with dissection time (model 16, χ^2^ = 14.843, p <0.01), but was independent of the antimalarial treatment (model 16, χ^2^ = 2.7322, p = 0.0983). In contrast, there was a very significant interaction between the SP-treatment and the time of dissection on the number of oocysts developing inside the mosquitoes (model 17, χ^2^ = 24.159, p < 0.01). Although the general trend was towards a decrease in the number of oocysts with time (Fig.1), mosquitoes having fed on a SP-treated bird had a consistently higher number of oocysts in their midgut than mosquitoes having fed on their control counterparts. These results are consistent across all the birds used in the experiment (Supplementary Materials, Fig S2-3). Fitting day as a continuous (rather than discrete) variable in the model revealed that the rate of decline of oocysts with time was significantly higher in control-fed mosquitoes (incidence rate ratio, IRR = 21%) than in SP-fed mosquitoes (IRR = 9.4%).

**Figure 1.**
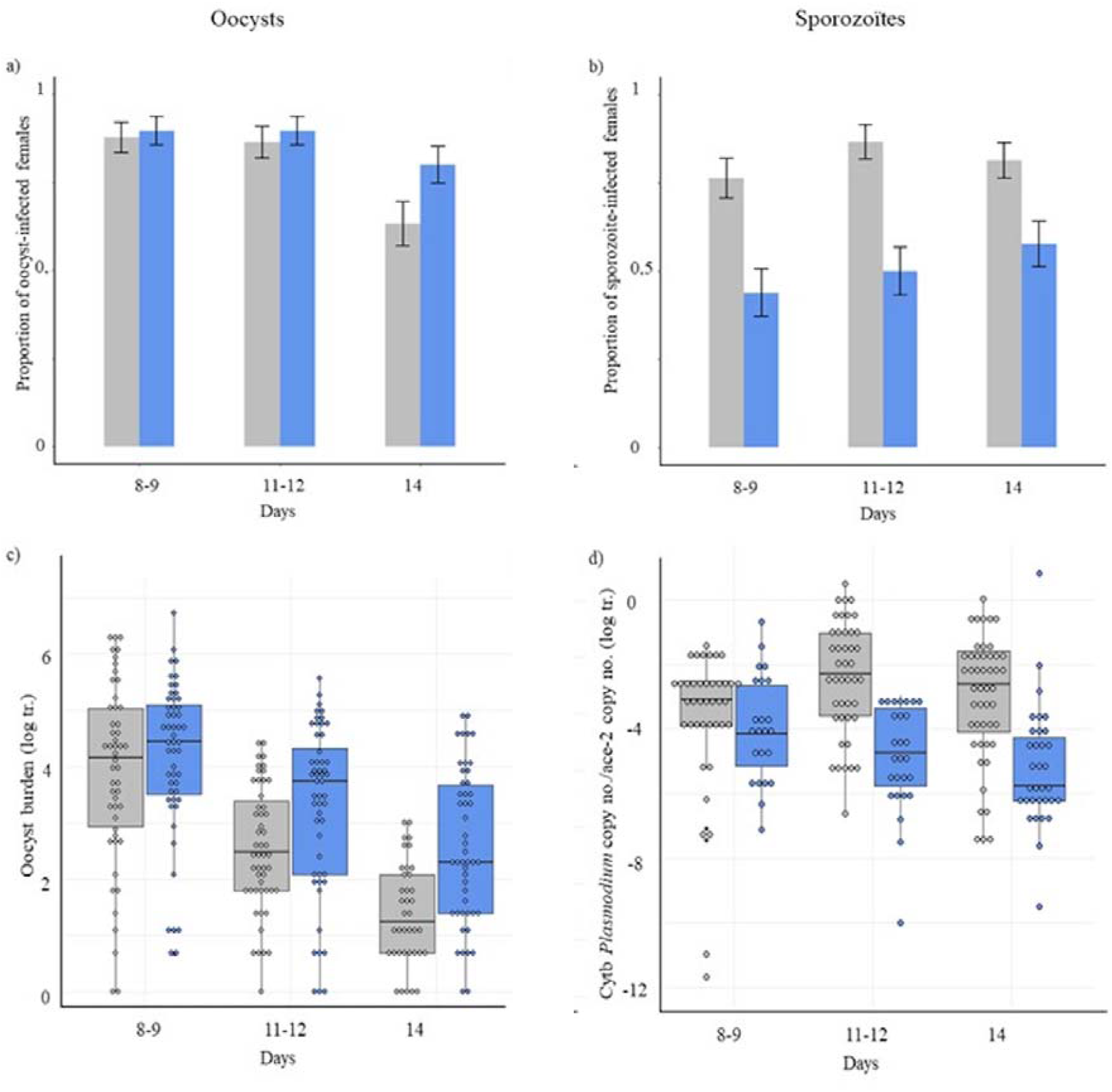
Prevalence and burden of oocysts and sporozoites in mosquitoes fed on a control-(grey) or SP-treated (blue) host for each sampling day (number of days post infection). a), b): oocyst and sporozoite prevalence, respectively. c), d): oocyst and sporozoite burden, respectively. Prevalence is represented as the mean ± standard error (calculated as sqrt(pq/n)). Burden is represented as a boxplot where with the median (horizontal lines), first and third quartiles (box above and below the medians). Vertical lines delimit 1.5 times the inter-quartile range above which individual counts are considered outliers and marked as circles.

Treatment had a significant effect on the prevalence of sporozoites within the mosquitoes (model 19, χ^2^ = 10.394, p <0.01). On average, parasites developing in mosquitoes having fed on an SP-treated host had a significantly lower probability of reaching the sporozoite stage than their control counterparts (55% vs 82%, respectively). Sporozoïte burden was also significantly lower in mosquitoes having fed on an SP-treated host, irrespective of the dissection date (model 20, *treatment*: χ^2^ = 9.8898, p < 0.01; *date*: χ^2^ = 3.1579, p= 0.2062; Fig. 1). As above, these results are consistent across all the birds used in the experiment (Supplementary Materials, Fig S2-3). Fitting day as a continuous (rather than discrete) variable in the model revealed that while in control-fed mosquitoes the number of sporozoites stayed roughly constant with time (slope not significantly different from 0, t= 1.66), in SP-treated mosquitoes, the number of sporozoites decreased significantly with time (t= 2.41).

#### 3.2.2. Artesunate experiment

Feeding on an AS-treated host had no impact on the prevalence of oocysts (model 22, χ^2^ = 0.854, p-value= 0.3554). Oocyst burden, on the other hand, showed the same pattern of decrease with time as in the previous experiment (model 23, χ^2^ = 211.91, p <0.01). There was a significant effect of treatment in interaction with the date of dissection (model 23, date*treatment: χ^2^ = 7.3787, p= 0.025). Post hoc analyses revealed the existence of a significant, albeit marginally, higher oocyst burden in treated hosts on days 11 and 12 (χ^2^ = 3.8886, p =0.0486) while no differences were observed in day 8,9 (χ^2^ = 0.0106, p =0.9179) and 14 (χ^2^ = 3.5452, p =0.0597).

Feeding on an AS-treated host, however, had no effect on either sporozoite prevalence (model 24: χ^2^ = 0.0106, p=0.9179), or burden (model 25, χ^2^ = 0.0002, p=0.9885, Fig 2

**Figure 2.**
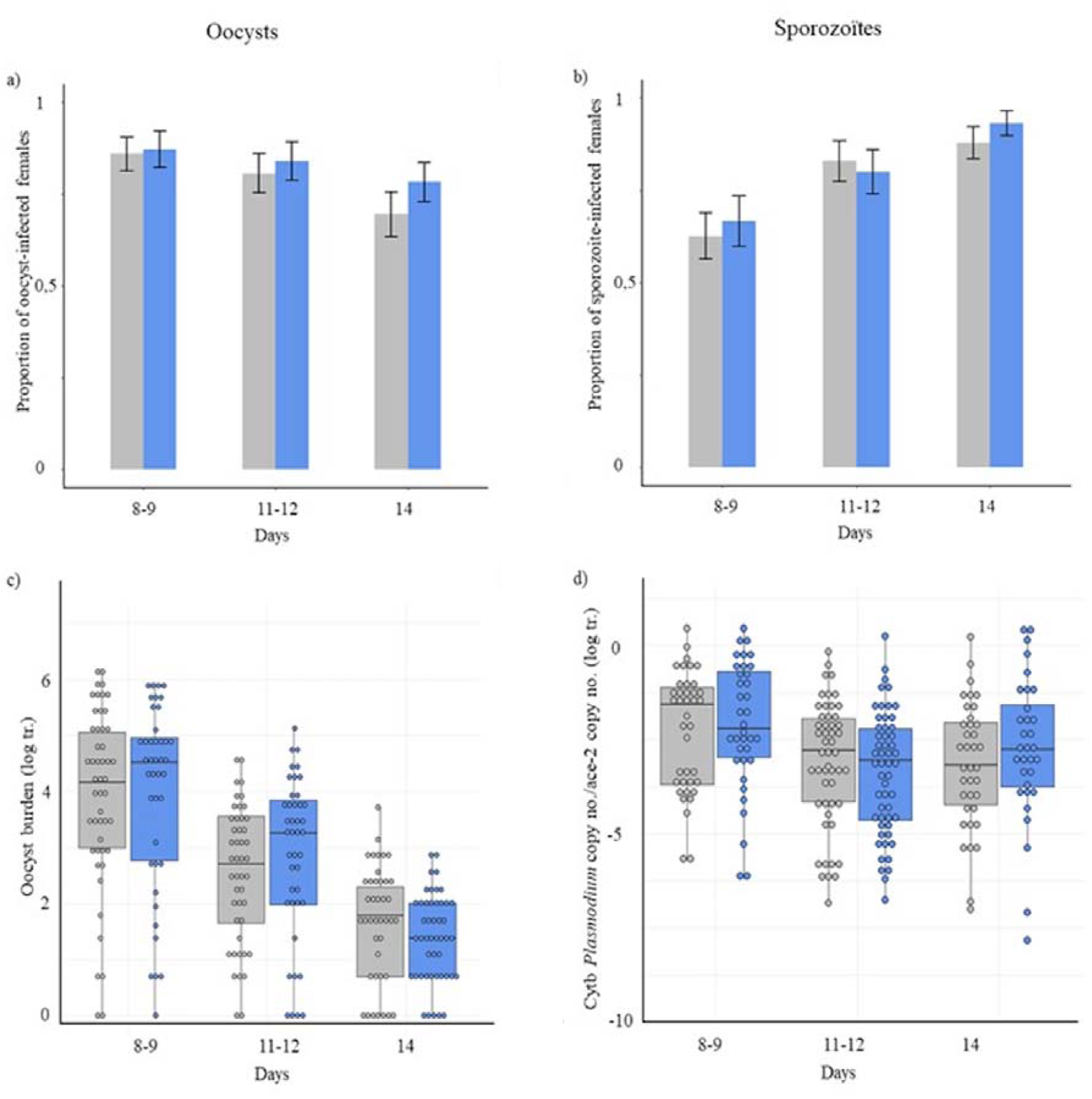
Prevalence and burden of oocysts and sporozoites in mosquitoes fed on a control-(grey) or AS-treated host (blue) for each sampling day (number of day post infection). a), b): oocyst and sporozoite prevalence, respectively. c), d): oocyst and sporozoite burden, respectively. Prevalence is represented as the mean ± standard error (calculated as sqrt(pq/n)). Burden is represented as a boxplot where with the median (horizontal lines), first and third quartiles (box above and below the medians). Vertical lines delimit 1.5 times the inter-quartile range above which individual counts are considered outliers and marked as circles.

## 4. Discussion

Artesunate and sulfadoxine-pyrimethamine are the cornerstone of modern antimalarial treatments in malaria-endemic areas. Millions of people across the world are treated every year with these drugs. Both antimalarials are extremely efficient at clearing the parasite from the red blood cells but, like most other drugs, they also come of a suite of adverse effects in humans (Medscape, 2020). The aim of our study was to establish whether this double toxicity, for both *Plasmodium* and its host, also takes place in the vector, thereby interfering on parasite transmission by mosquitoes. More precisely we aimed to establish: 1) whether mosquitoes feeding on an AS or SP treated host suffer any adverse fitness effects from the drugs, and 2) whether the drugs are toxic for the oocysts and sporozoites developing inside a mosquito

For this purpose, we carried out several factorial experiments feeding both uninfected mosquitoes and mosquitoes with a 6-day old infection (corresponding to the early stages of oocyst formation in *P. relictum*, Pigeault, 2015) on drug treated (AS or SP) and control hosts. We then quantified the life history traits of the mosquito (fecundity, longevity) and the oocyst (midgut) and sporozoite (salivary gland) stages of the parasites developing inside them.

Our results show what seem to be mostly minor effects of the drugs on the life history traits of mosquitoes feeding from a treated host. Amongst the two life history traits quantified that are known to be key for malaria transmission: mosquito longevity (Smith and McKenzie, 2004) and host feeding probability (Cornet et al., 2019), neither were found to be affected by the drug treatments. Previous work on the longevity effects of drugs has shown that *An. gambiae* mosquitoes membrane-fed on a gametocyte culture containing high concentrations SP had significantly shorter lifespans (Kone et al., 2010). Whether this is due to differences in the experimental system or, more likely, to key differences in experimental conditions (Kone et al added a high SP dose to a gametocyte culture) is unclear. In our experiments, some significant interactions were, however, found that may be worthy of further study. Females that fed on an SP-treated bird laid eggs on average 8 hours earlier than those fed on control birds, a result that agrees with previous studies showing that *Culex pipiens* mosquitoes are able to advance their oviposition schedule when faced with adverse conditions (Vézilier et al., 2015). In addition, mosquitoes taking small blood meals from AS-treated birds saw their probability of laying an egg raft reduced by 37% as compared to their control counterparts. In humans, artesunate use is frequently associated with haemolitic anaemia as evidenced by a decline in blood haemoglobin levels and an increase in reticulocyte counts (Burri et al., 2014; Sowunmi et al., 2017). Had a similar phenomenon taken place in our birds, mosquitoes taking a small blood meal from AS-treated hosts would not have obtained enough haemoglobin to produce a batch of eggs (Ferguson et al., 2003; Vézilier et al., 2012; Zhou et al., 2007). We found no difference in the total number of red blood cells between AS-treated and untreated birds, but since our analysis did not allow us to distinguish between young (reticulocyte) and mature red blood cells, we could not establish whether artesunate induces anaemia in this system.

In contrast to the effects observed on mosquito life history traits which, interesting as they may be from a biological standpoint are unlikely to bear significant consequences for the epidemiology of the disease, the substantial reduction in both sporozoite prevalence (−30%) and burden (−80%) in mosquitoes having taken an SP-treated blood meal, may result in a drastic reduction in the transmission potential of the parasite. Sulfadoxine and pyrimethamine act synergistically to inhibit the activity of dihydropteroate synthase (DHPS) and dihydrofolate reductase (DHFR), respectively, thus inhibiting the folic acid metabolism of the parasite (Hopkins Sibley et al., 2001). Folic acid is vital for the biosynthesis of purines and pyrimidines, which are essential for DNA synthesis and cell multiplication (Kirk et al., 1976). The mitotic-blocking properties of pyrimethamine were first gleaned through work done on *Plasmodium gallinaceum* were birds treated with high concentrations of pyrimethamine showed arrested schizont division and fewer merozoites were produced (Aikawa and Beaudoin, 1968). Since then, the schistocidal effect of pyrimethamine has been confirmed in several other systems (Delves et al., 2012; Vincke, 1970). In contrast, work on the effect of pyrimethamine on *Plasmodium* sporogony in the mosquito has produced contrasting results. The overwhelming majority of these studies tested the so-called *prophylactic* effect of pyrimethamine on the mosquito, that is, the effect of the drug when administered prior to or concomitantly with the infected blood meal (Table 1). These studies found that when administered with the infected blood meal, pyrimethamine averted the arrival of sporozoites to the salivary gland. There was, however, no consensus on the mechanisms underlying this sporozoite-inhibitory effect: pyrimethamine may have rendered gametocytes uninfective (Foy and Kondi, 1952), prevented the ookinetes from traversing the midgut wall (Bray et al., 1959), or prevented the oocysts from reaching maturity (Terzian, 1970; Terzian et al., 1968). More recent work seems to confirm that pyrimethamine in combination with sulfadoxine, decreases the infectiousness of gametocytes (Beavogui et al., 2010; Kone et al., 2010) and Delves et al. have reported that pyrimethamine and other antifolates result in a strong (> 90%) inhibition of male gametocyte exflagellation, while having virtually no effect on female gametocytes (Delves et al., 2012, 2013) thus effectively strongly skewing the parasites’ operational sex ratio. These studies collectively suggest that a prophylactic administration of SP has transmission-blocking effect through the inhibition of the early (gametocyte) stages within the mosquito.

**Table 1:** Summary of the main studies investigating the inhibitory effect of pyrimethamine (PYR) alone or in combination with sulfadoxine (SFX) for oocyst and sporozoite formation. In *prophylactic* protocols the drug is administered before or concomitantly with the infected blood meal (mosquitoes ingest the drug at the same time as the infective gametocytes). In *curative* protocols, the mosquito is first infected and then provided with a second blood meal containing the drug.

| *Plasmodium* species | Mosquito species | Dose* | Oocyst inhibition | Sporozoite inhibition | Reference |
| --- | --- | --- | --- | --- | --- |
| ***Prophylactic administration*** |  |  |  |  |  |
| *P. falciparum* | *An. gambiae* | 20 mg PYR /ind | - | YES | (Foy and Kondi, 1952) |
| *P. falciparum* | *An. stephensi* | 25 mg PYR /ind | YES<sup>1</sup> | NO<sup>2</sup> | (Shute and Maryion, 1954) |
| *P. falciparum* | *An. gambiae*<br>*An. melas* | 25-50 mg PYR /ind | YES | YES | (Bray et al., 1959) |
| *P. falciparum* | *An. quadrimaculatus*<br>*An. freeborni* | 50-100 mg PYR /ind | YES<sup>1</sup> | YES | (Burgess and Young, 1959) |
| *P. falciparum* | *An. gambiae* | 12-50 mg PYR/ind | YES<sup>1,3</sup> | YES<sup>2,3</sup> | (Gunders, 1961) |
| *P. falciparum* | *An. stephensi* | 0.00001% PYR (ss) | - | YES<sup>2</sup> | (Gerberg, 1971) |
| *P. falciparum* | *An. stephensi* | 10<sup>-7</sup> M PYR (mf) | YES<sup>1</sup> | - | (Chutmongkonkul et al., 1992) |
| *P. falciparum* | *An. gambiae* | 75 mg PYR<br>1500 mg SFX | YES<sup>1</sup> | - | (Hogh et al., 1998) |
| *P. falciparum* | *An. gambiae* | 1.25 mg/kg PYR<br>25 mg/kg SFX | YES | - | (Beavogui et al., 2010) |
| *P. falciparum* | *An. arabiensis* | 25 mg/kg SFX<br>1.25 mg/kg PYR | YES<sup>1,4</sup> | - | (Robert et al., 2000) |
| *P. vivax* | *An. stephensi* | 50 mg PYR /ind | YES<sup>1</sup> | NO<sup>2</sup> | (Shute and Maryion, 1954) |
| *P. vivax* | *An. stephensi* | 0.002 gr PYR /ml (ss) | YES<sup>1</sup> | YES | (Terzian et al., 1968) |
| *P. cynomolgui* | *An. stephensi* | 0.001 gr PYR /ml (ss) | YES<sup>1</sup> | YES | (Terzian, 1970) |
| *P. cynomolgui* | *An. stephensi* | 0.00001% PYR (ss) | - | YES<sup>2</sup> | (Gerberg, 1971) |
| *P. cynomolgui* | *An. maculatus* | 3 mg PYR /kg | YES | YES | (Omar et al., 1973) |
| *P. berghei* | *An. stephensi* | 2.5 - 20mg PYR /kg | YES | YES | (Vincke, 1970) |
| *P. berghei* | *An. stephensi* | 20mg PYR /kg | YES | YES | (Shinondo et al., 1994) |
| *P. berghei* | *An. stephensi* | Serum from PYR/SFX treated patients (mf) | YES<sup>1</sup> | - | (Hogh et al., 1998) |
| *P. gallinaceum* | *Ae. aegypti* | 0.028 mg/kg PYR<br>210 mg/kg SFX | - | YES | (Ramakrishnan et al., 1963) |
| *P. gallinaceum* | *Ae. aegypti* | 0.001% and<br>0.0001% PYR (ss) | YES<sup>1</sup> | YES<sup>2</sup> | (Terzakis, 1971) |
| *P. gallinaceum* | *Ae. aegypti* | 0.00001% PYR (ss) | - | YES<sup>2</sup> | (Gerberg, 1971) |
| ***Curative administration*** |  |  |  |  |  |
| *P. falciparum* | *An. gambiae*, *An. melas* | 25-50 mg PYR<br>4 days post infection | YES<sup>1</sup> | NO<sup>2</sup> | (Bray et al., 1959) |
| *P. falciparum* | *An. gambiae* | 1μM PYR 2-4 days post-infection (mf) | YES<sup>1</sup> | NO<sup>2</sup> | (Teklehaimanot et al., 1985) |
| *P. falciparum* | *An. stephensi* | 10<sup>-7</sup> M PYR 4 days post infection (mf) | NO | - | (Chutmongkonkul et al., 1992) |

\* Drug administered directly to host unless otherwise stated: (ss): drug administered in sugar solution, (mf): drug added to blood in a membrane feeder. <sup>1</sup> Inhibition was partial (some oocysts present); <sup>2</sup> Sporozoites were observed but not quantified; <sup>3</sup> No untreated controls; <sup>4</sup> Chloroquine-treated patients used as a control.

Our experiments were carried out using a *curative* protocol, i.e. the drug was administered to mosquitoes carrying a 6-day old *Plasmodium* infection which, in this system, corresponds to the initial stages of the oocyst invasion of the midgut. The drastic decrease obtained in both sporozoite prevalence (Fig 1b) and burden (Fig 1d) demonstrate that SP has an additional effect on parasite development, which is downstream from its toxicity to (male) gametes. Although the underlying mechanism remains to be established, these results are consistent with the mitosis-blocking properties of antifolates observed in the blood stages of parasites, which may here have prevented the multiple rounds cell division that take place inside the syncytial oocyst prior to the liberation of the sporozoites (Gerald et al., 2011). Recent experiments using luciferase-expressing *Plasmodium berghei* parasites cultured *in vitro* (Azevedo et al., 2017) have observed a significant reduction in the luminescence of oocysts after adding 10µM pyrimethamine to the parasite culture. As the luciferase was under the control of the parasite’s circumsporozoite protein (*Pb*CSP) promoter regions, a reduction in the number of sporozoites produced inside the oocyst therefore seems like a plausible explanation for the observed reduction in bioluminescence.

The strong reduction in sporozoite prevalence and burden in SP-fed mosquitoes is all the more notable for being associated with a significant concomitant increase in the number of oocysts in the midgut (Fig 1c). One potential explanation is that the powerful antibiotic properties of sulfadoxine and pyrimethamine may have altered the microbiota of the mosquito midgut (Capan et al., 2010) which has been shown to be correlated with oocyst development (Gendrin et al., 2015; Saraiva et al., 2016). We are not aware of any study that has investigated the antibiotic properties of SP against the mosquito midgut flora. To our knowledge, this is the first time that such an increase in oocysts following a drug-treated blood meal has been reported in any study, which raises some interesting questions regarding the timing of SP administration with respect to the arrival of ookinetes to the midgut wall. In addition, we observed an interesting pattern whereby the decline in oocyst numbers with time, a natural process that takes place as mature oocysts burst to produce sporozoites, happens more rapidly in control-fed than in SP-fed mosquitoes, which may be indicative of a delay in oocyst development in the latter. Other possibilities for the increased oocystaemia observed, such as a SP-induced immunosuppression or SP-induced facilitation of the ookinete midgut invasion would also merit further study. Irrespective of the underlying mechanism, our results indicate that SP exerts two opposing effects on the parasite’s sporogonic development within the mosquito: one that facilitates the midgut invasion of oocysts, followed by another that blocks the production of sporozoites within them.

In contrast to SP, AS treatment had no discernible effects on sporozoite burden and only a minor effect, detectable only 11-12 post infection, on the number of oocysts. These results are in agreement with previous work showing that artesunate and other artemisinine derivatives have a considerable gametocytocidal effect in humans but no effect on the mosquito stages of the parasite (Butcher, 1997; Wadi et al., 2019).

We are acutely aware that the effects of curative administration of SP on oocyst and sporozoite burden may not directly translatable to human malaria. SP is widely used as a preventive treatment for uninfected children (SMC) and pregnant women (IPT), with millions of doses being provided every year across the African continent (Van Eijk et al., 2011). The transmission-blocking effect of a curative administration of SP demonstrated here would be relevant when infected mosquitoes bite these SP-treated individuals (in the field, mosquitoes go through several gonotrophic, bloodmeal - egg laying - bloodmeal, cycles, Bomblies, 2014). To confirm the curative effect of SP in human malaria infections, experiments where infected mosquitoes are membrane-fed on treated uninfected blood could be carried out, with the caveat that membrane feeding and direct feeding on human volunteers may render different results (Beavogui et al., 2010; Butcher, 1989; Wadi et al., 2018). Provided the results obtained here are repeatable in human malaria, the epidemiological and evolutionary consequences of the preventive use of SP in malaria-endemic countries could be substantial. Fewer sporozoite-carrying mosquitoes (−30%), and fewer sporozoites in the salivary gland (−80%) should translate into lower transmission rates, even accounting for a non-linear correlation between sporozoite load and transmission (Aleshnick et al., 2019). The evolutionary effects may not be less important. Current work largely assumes that the strongest selective pressures for drug resistance operate on the treated host. As these results show, the strong bottleneck for sporozoites in the mosquito may act as an additional selective pressure which may help maintain drug resistance in the field even when the drug is not used to treat infected hosts, as is the case in the current mass administration of pyrimethamine for ITP and SMC. More generally, these results also highlight the need for further studies on the effects of the transmission-blocking compounds on each stage of the parasite’s cycle within the mosquito. The results of standard membrane feeding assays (SMFAs), considered to be the gold standard for assessing the efficiency of transmission blocking interventions, are reported as a percent reduction in the number of oocysts compared to a control (Nunes et al., 2014; Paton et al., 2019), with current efficacy thresholds set at around a 80% reduction. As shown here, drugs can have contrasting effects on different stages of the parasite’s sporogonic cycle highlighting the potential drawbacks of assessing drug-based transmission-blocking interventions based on oocyst quantifications alone.

## 5. Acknowledgements

We would like to thank Tanguy Lagache for his help with the experiments. This work was funded through the ANR-16-CE35-0001-01 (‘EVODRUG’).

## Supplementary Materials

The transmission-blocking effects of antimalarial drugs revisited: fitness costs and sporontocidal effects (Villa et al)

**Table S1.** Description of the statistical models used to analyze the impact of drugs on mosquito life history traits. Models with binomial error structure require a concatenated response variable binding together the number of successes and failures for a given outcome. N gives the number of mosquitoes included in each analysis. “Maximal model” represents the complete set of explanatory variables (and their interactions) included in the model. “Minimal model” represents the model containing only the significant variables and their interactions. Round brackets indicate that the variable was fitted as a random factor. Square brackets indicate the error structure used (n: normal errors, b: binomial errors). date: sampling day, status: alive/dead on sampling day, fed/unfed: number of fed/unfed mosquitoes, hm: haematin excreted (proxi for blood meal size), plt: plate used for the colorimetric quantification haematin, hmr: residuals of hm by plate, eggs: number of eggs laid, inf: mosquito infection status (infected/uninfected), TR: mosquito fed on treated/untreated bird.

| Variable of interest |  | Response variable | Model Nb. | N | Maximal model | Minimal model | R subroutine [err struct.] |
| --- | --- | --- | --- | --- | --- | --- | --- |
| *Effect of AS on mosquito traits* |  |  |  |  |  |  |  |
| RBC | RBC/ml | RBC | 5 | 5 | TR | 1 | K-W |
| Survival | Overall survival | (date, status) | 14 | 414 | TR*inf + (1\|bird) | (1\|bird) | coxme |
|  | % mosquitoes surviving until day 14 | cbind (dead, alive) | 15 | 10 | TR*inf + (1\|bird) | (1\|bird) | lmer [b] |
| Blood meal size | Blood-fed | cbind (fed, unfed) | 9 | 966 | TR*inf + (1\|bird) | (1\|bird) | lmer [b] |
|  | Blood meal size | hmr (lm (hm ~ plt) | 10 | 414 | TR*inf + (1\|bird) | (1\|bird) | lmer [n] |
| Fecundity | Egg laying probability | cbind (laid, not laid) | 13 | 440 | hmr*TR*inf + (1\|bird) | TR*hmr + inf + (1\|bird) | lmer [b] |
|  | Oviposition day | day | 11 | 337 | hmr*TR*inf | hmr | clm |
|  | Number of eggs per raft | eggs | 12 | 337 | hmr*TR + inf*day + (1\|bird) | hmr + (1\|bird) | glmmTMB |
| *Effect of SP on mosquito traits* |  |  |  |  |  |  |  |
| RBC | RBC/ml | RBC | 3 | 6 | TR | 1 | K-W |
| Survival | overall survival | (date, statut) | 7 | 564 | TR*inf + (1\|bird) | inf + (1\|bird) | coxme |
|  | % mosquitoes surviving until day 14 | cbind(dead, alive) | 8 | 12 | TR*inf + (1\|bird) | (1\|bird) | lmer [b] |
| Blood meal | Blood-fed | cbind (fed, unfed) | 1 | 1045 | TR*inf + (1\|bird) | (1\|bird) | lmer [b] |
|  | Blood meal size | hmres (lm (hm ~ Plq) | 2 | 454 | TR*inf + (1\|bird) | (1\|bird) | lmer [n] |
| Fecundity | Egg-laying probability | cbind (laid, not laid) | 4 | 378 | hmr*TR*inf + (1\|bird) | TR*inf + (1\|bird) | lmer [b] |
|  | Oviposition day | day | 5 | 312 | hmr*TR*inf | hmr*inf + TR | clm |
|  | Number of eggs per raft | eggs | 6 | 312 | hmr*TR + inf*day + (1\|bird) | day + (1\|bird) | glmmTMB |

**Table S2.**
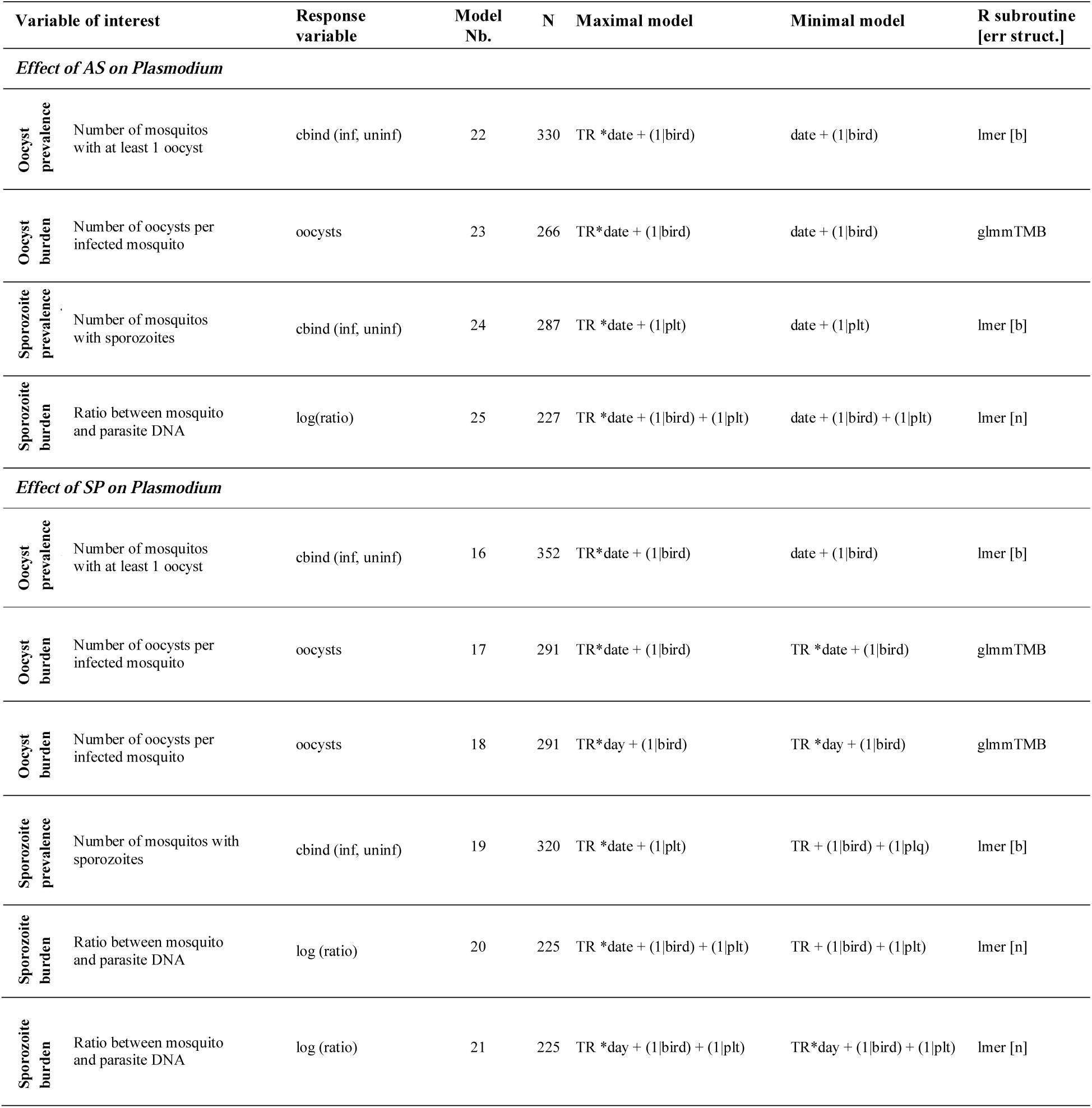
Description of the statistical models used to analyze the impact of drugs on *Plasmodium* prevalence and burden. Models with binomial error structure require a concatenated response variable binding together the number of successes and failures for a given outcome. N gives the number of mosquitoes included in each analysis. “Maximal model” represents the complete set of explanatory variables (and their interactions) included in the model. “Minimal model” represents the model containing only the significant variables and their interactions. Round brackets indicate that the variable was fitted as a random factor. Square brackets indicate the error structure used (n: normal errors, b: binomial errors). date: mosquito dissection day (discrete variable), day: mosquito dissection day (continuous variable), inf: mosquito infection status (infected/uninfected), plt: plate used for the sporozoite quantification (qPCR), TR: mosquito fed on treated/untreated bird.

**Figure S1.**
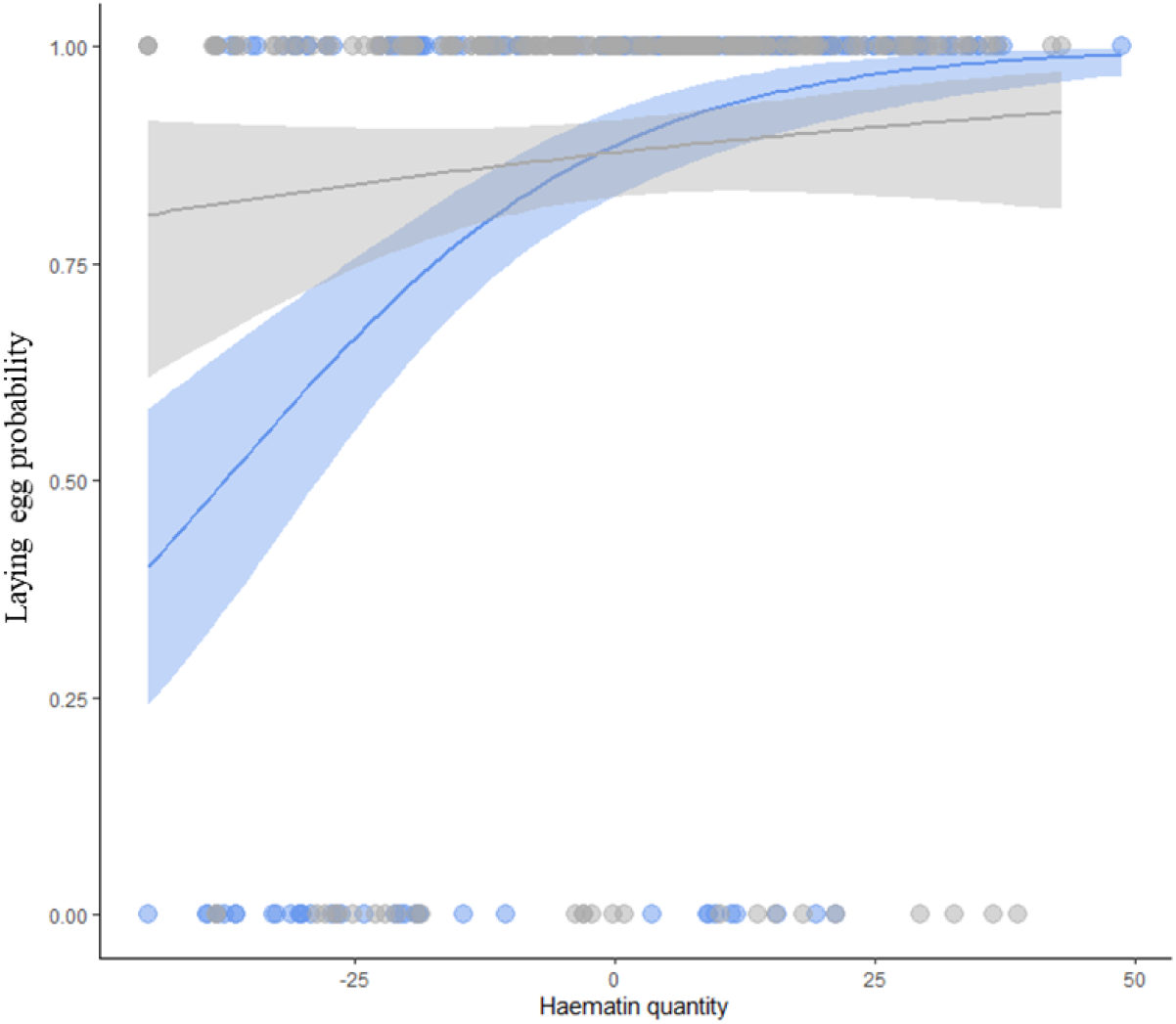
Laying egg probability as a function of the haematine excreted (represented here by the residuals of a model containing ‘plate’ as an explanatory variable, see materials and methods.) Blue: mosquitoes fed on an AS-treated bird, grey: mosquitoes fed on a control bird. Each point represents an individual, lines are fitted using a logistic regression the grey areas are the 95% confident intervals.

**Figure S2.**
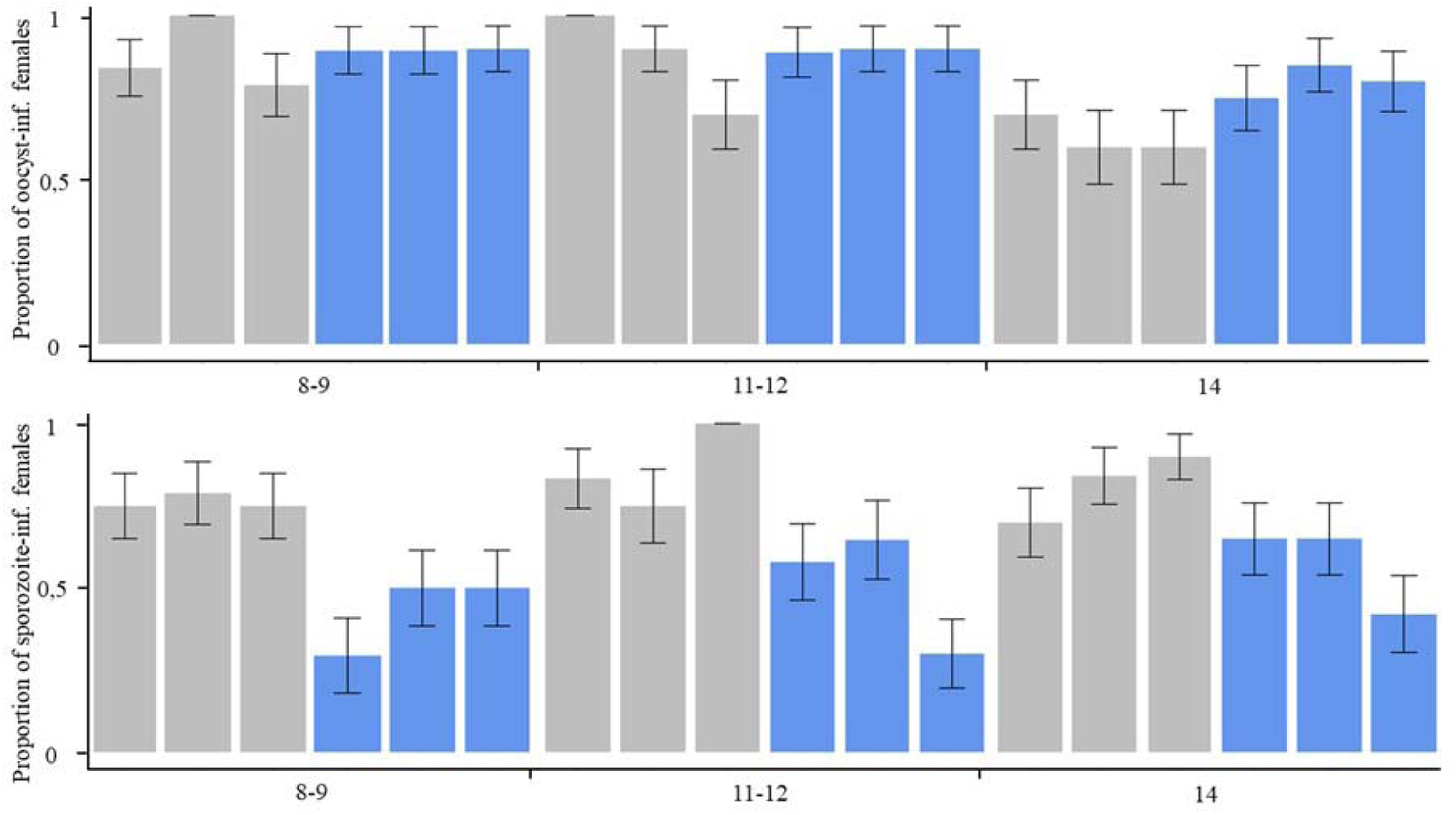
Prevalence of oocysts (top) and sporozoites (bottom) in mosquitoes fed on each of the 3 control (grey) and SP-treated (blue) birds, for each of the 3 sampling days (8-9, 11-12 and 14). Prevalence is represented as the mean ± standard error (calculated as sqrt(pq/n)). Burden is represented as a boxplot where with the median (horizontal lines), first and third quartiles (box above and below the medians). Vertical lines delimit 1.5 times the inter-quartile range above which individual counts are considered outliers and marked as circles.

**Figure S3.**
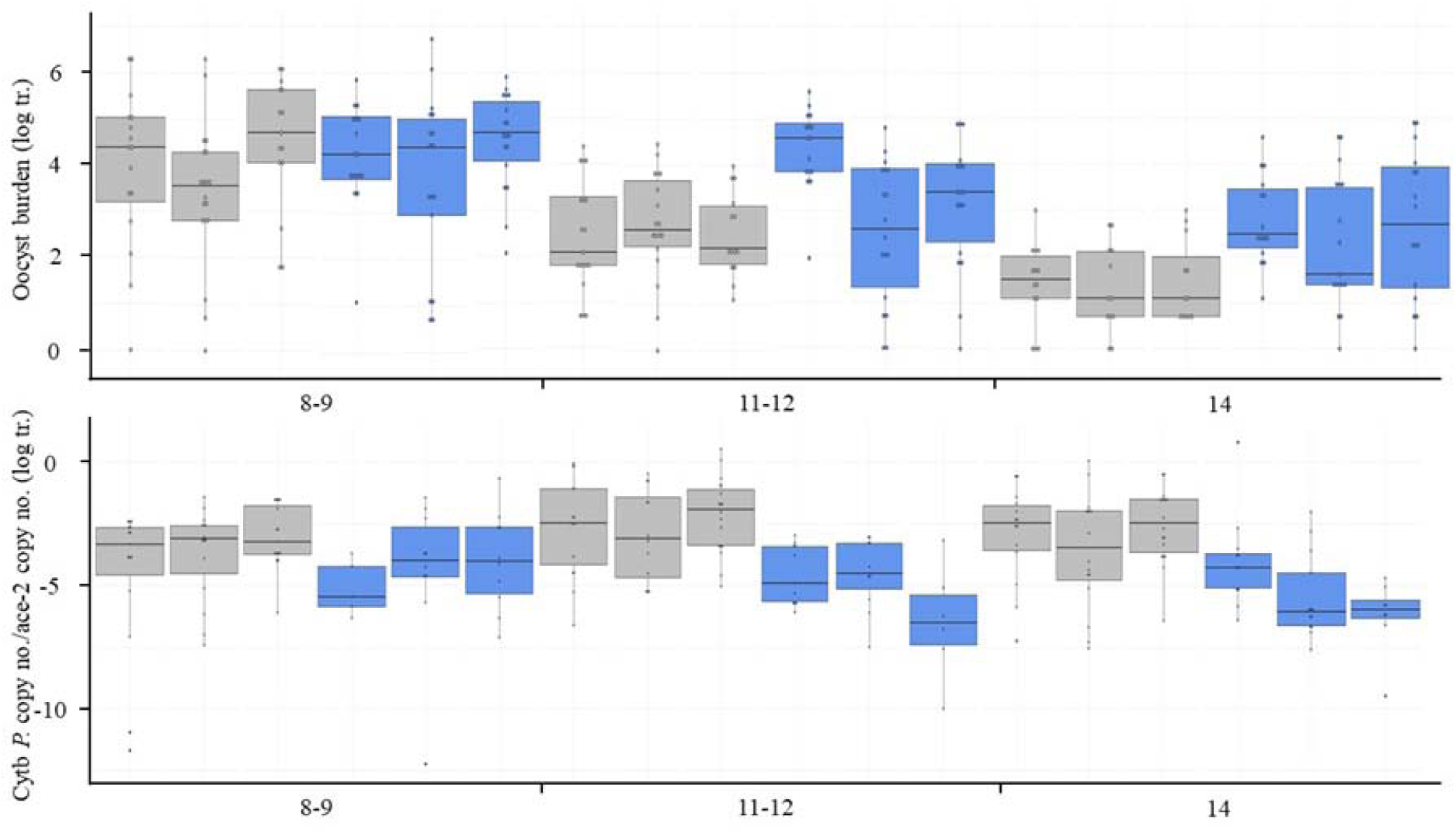
Oocyst (top) and sporozoite (bottom) burden in mosquitoes fed on each of the 3 control (grey) and SP-treated (blue) birds, for each of the 3 sampling days (8-9, 11-12 and 14). Burden is represented as a boxplot where with the median (horizontal lines), first and third quartiles (box above and below the medians). Vertical lines delimit 1.5 times the inter-quartile range above which individual counts are considered outliers and marked as circles.

## Notes

Declaration of interests: none

### Competing Interest Statement

The authors have declared no competing interest.

## References

Aikawa, M., Beaudoin, R.L., 1968. Studies on nuclear division of a malarial parasite under pyrimethamine treatment. J. Cell Biol. 39, 749–754. doi: 10.1083/jcb.39.3.749

Aleshnick, M., Ganusov, V. V., Nasir, G., Yenokyan, G., Sinnis, P., 2019. Experimental determination of the force of malaria infection reveals a non-linear relationship to mosquito sporozoite loads. bioRxiv 1–40. doi: 10.1101/830299

Azevedo, R., Markovic, M., Machado, M., Franke-Fayard, B., Mendes, A.M., Prudêncio, M., 2017. Bioluminescence method for in vitro screening of *Plasmodium* transmission-blocking compounds. Antimicrob Agents Chemother 61, e02699–16.

Balint, G.A., 2001. Artemisinin and its derivatives: an important new class of antimalarial agents. Pharmacol. Ther. 90, 261–265. doi: 10.1016/S0163-7258(01)00140-1

Beavogui, A.H., Djimde, A.A., Gregson, A., Toure, A.M., Dao, A., Coulibaly, B., Ouologuem, D., Fofana, B., Sacko, A., Tekete, M., Kone, A., Niare, O., Wele, M., Plowe, C. V., Picot, S., Doumbo, O.K., 2010. Low infectivity of *Plasmodium falciparum* gametocytes to *Anopheles gambiae* following treatment with sulfadoxine-pyrimethamine in Mali. Int. J. Parasitol. 40, 1213–1220. doi: 10.1016/j.ijpara.2010.04.010

Blagborough, A.M., Churcher, T.S., Upton, L.M., Ghani, A.C., Gething, P.W., Sinden, R.E., 2013. Transmission-blocking interventions eliminate malaria from laboratory populations. Nat. Commun. 4, 1–7. doi: 10.1038/ncomms2840

Bolker, B.M., 2008. Ecological models and data in R, Ecological Models and Data in R. doi: 10.1111/j.1442-9993.2010.02210.x

Bomblies, A., 2014. Agent-based modeling of malaria vectors: the importance of spatial simulation. Parasites and Vectors 7, 308. doi: 10.1186/1756-3305-7-308

Bray, R.S., Burgess, R.W., Fox, R.M., Miller, M.J., 1959. Effect of pyrimethamine upon sporogony and pre-erythrocytic schizogony of *Laverania falciparum*. Bull. World Health Organ. 21, 233–238.

Buckling, A., Crooks, L., Read, A., 1999. *Plasmodium chabaudi* : effect of antimalarial drugs on gametocytogenesis. Exp. Parasitol. 93, 45–54. doi: 10.1006/EXPR.1999.4429

Burgess, R.W., Young, M.D., 1959. The development of pyrimethamine resistance by *Plasmodium falciparum*. Bull. World Health Organ. 20, 37–46.

Burri, C., Ferrari, G., Ntuku, H.M., Kitoto, A.T., Duparc, S., Hugo, P., Mitembo, D.K., Lengeler, C., 2014. Short report: delayed anemia after treatment with injectable artesunate in the Democratic Republic of the Congo: a manageable issue. Am. J. Trop. Med. Hyg. 91, 821–823. doi: 10.4269/ajtmh.14-0149

Butcher, G., 1997. Antimalarial drugs and the mosquito transmission of *Plasmodium*. Int. J. Parasitol. 27, 975–987.

Butcher, P., 1989. Mechanisms of immunity to malaria and the possibilities of a blood-stage vaccine: a critical appraisal. Parasitology 98, 315–327. doi: 10.1017/S0031182000062247

Capan, M., Mombo-Ngoma, G., Makristathis, A., Ramharter, M., 2010. Anti-bacterial activity of intermittent preventive treatment of malaria in pregnancy: comparative *in vitro* study of sulfadoxine-pyrimethamine, mefloquine, and azithromycin. Malar. J. 9, 303. doi: 10.1186/1475-2875-9-303

Chotivanich, K., Sattabongkot, J., Udomsangpetch, R., Looareesuwan, S., Day, N.P.J., Coleman, R.E., White, N.J., 2006. Transmission-blocking activities of quinine, primaquine, and artesunate. Antimicrob. Agents Chemother. 50, 1927–1930. doi: 10.1128/AAC.01472-05

Churcher, T.S., Bousema, T., Walker, M., Drakeley, C., Schneider, P., Ouédraogo, A.L., Basáñez, M.G., 2013. Predicting mosquito infection from *Plasmodium falciparum* gametocyte density and estimating the reservoir of infection. Elife e00626. doi: 10.7554/eLife.00626

Chutmongkonkul, M., Maier, W.A., Seitz, H.M., 1992. *Plasmodium falciparum* : effect of chloroquine, halofantrine and pyrimethamine on the infectivity of gametocytes for *Anopheles stephensi* mosquitoes. Ann. Trop. Med. Parasitol. 86, 103–110. doi: 10.1080/00034983.1992.11812639

Coatney, G.R., Cooper, W.C., Eddy, N.B., Greenberg, J., 1953. Survey of antimalarial agents: chemotherapy of *Plasmodium gallinaceum* infections; toxicity; correlation of structure and action. Public Heal. Serv. Publ.

Cohuet, A., Osta, M.A., Morlais, I., Awono-Ambene, P.H., Michel, K., Simard, F., Christophides, G.K., Fontenille, D., Kafatos, F.C., 2006. *Anopheles* and *Plasmodium* : from laboratory models to natural systems in the field. EMBO Rep. 7, 1285–1289. doi: 10.1038/sj.embor.7400831

Cornet, S., Nicot, A., Rivero, A., Cator, L., 2019. Avian malaria alters blood feeding of *Culex pipiens* mosquitoes. Malar. J. 18, 82.

Cornet, S., Nicot, A., Rivero, A., Gandon, S., 2014. Evolution of plastic transmission strategies in avian malaria. PLoS Pathog. 10. doi: 10.1371/journal.ppat.1004308

Crawley, M.J., 2007. The R book, Imperial C. ed. UK.

Delves, M., Plouffe, D., Scheurer, C., Meister, S., Wittlin, S., Winzeler, E.A., Sinden, R.E., Leroy, D., 2012. The activities of current antimalarial drugs on the life cycle stages of *Plasmodium* : a comparative study with human and rodent parasites. PLoS Med. 9. doi: 10.1371/journal.pmed.1001169

Delves, M.J., Miguel-Blanco, C., Matthews, H., Molina, I., Ruecker, A., Yahiya, S., Straschil, U., Abraham, M., León, M.L., Fischer, O.J., Rueda-Zubiaurre, A., Brandt, J.R., Cortés, Á., Barnard, A., Fuchter, M.J., Calderón, F., Winzeler, E.A., Sinden, R.E., Herreros, E., Gamo, F.J., Baum, J., 2018. A high throughput screen for next-generation leads targeting malaria parasite transmission. Nat. Commun. 9, 1–13. doi: 10.1038/s41467-018-05777-2

Delves, M.J., Ruecker, A., Straschil, U., Lelièvre, J., Marques, S., López-Barragán, M.J., Herreros, E., Sinden, R.E., 2013. Male and female *Plasmodium falciparum* mature gametocytes show different responses to antimalarial drugs. Antimicrob. Agents Chemother. 57, 3268–3274. doi: 10.1128/AAC.00325-13

Dong, Y., Aguilar, R., Xi, Z., Warr, E., Mongin, E., Dimopoulos, G., 2006. *Anopheles gambiae* immune responses to human and rodent *Plasmodium* parasite species. PLoS Pathog. 2, e52. doi: 10.1371/journal.ppat.0020052

Ferguson, H.M., Rivero, A., Read, A.F., 2003. The influence of malaria parasite genetic diversity and anaemia on mosquito feeding and fecundity. Parasitology 127, 9–19. doi: 10.1017/S0031182003003287

Foy, H., Kondi, A., 1952. Effect of daraprim on the gametocytes of *Plasmodium falciparum*. Trans. R. Soc. Trop. Med. Hyg. doi: 10.1016/0035-9203(52)90084-9

Gendrin, M., Rodgers, F.H., Yerbanga, R.S., Ouédraogo, J.B., Basáñez, M.G., Cohuet, A., Christophides, G.K., 2015. Antibiotics in ingested human blood affect the mosquito microbiota and capacity to transmit malaria. Nat. Commun. 6, 1–7. doi: 10.1038/ncomms6921

Gerald, N., Mahajan, B., Kumar, S., 2011. Mitosis in the human malaria parasite *Plasmodium falciparum*. Eukaryot. Cell 10, 474–482. doi: 10.1128/EC.00314-10

Gerberg, E.J., 1971. Evaluation of antimalarial compounds in mosquito test systems. Trans. R. Soc. Trop. Med. Hyg. 65, 358–363.

Gunders, A.E., 1961. The effect of a single dose of pyrimethamine and primaquine in combination upon gametocytes and sporogony of *Laverania falcipara* (=*Plasmodium falciparum*) in Liberia. Bull. World Health Organ. 24, 650–653.

Hewitt, R., 1940. Bird Malaria, The american journal of hygiene.

Hogh, B., Gamage-Mendis, A., Butcher, G.A., Thompson, R., Begtrup, K., Mendis, C., Enosse, S.M., Dgedge, M., Barreto, J., Eling, W., Sinden, R.E., 1998. The differing impact of chloroquine and pyrimethamine/sulfadoxine upon the infectivity of malaria species to the mosquito vector. Am. J. Trop. Med. Hyg. 58, 176–182. doi: 10.4269/ajtmh.1998.58.176

Hopkins Sibley, C., Hyde, J.E., Sims, P.F.G., Plowe, C. V., Kublin, J.G., Mberu, E.K., Cowman, A.F., Winstanley, P.A., Watkins, W.M., Nzila, A.M., 2001. Pyrimethamine-sulfadoxine resistance in *Plasmodium falciparum* : what’s next? TRENDS Parasitol. 17, 582–588.

Hou, L., Huang, H., 2016. Immune suppressive properties of artemisinin family drugs. Pharmacol. Ther. 166, 123–127. doi: 10.1016/j.pharmthera.2016.07.002

Karunajeewa, H.A., Salman, S., Mueller, I., Baiwog, F., Gomorrai, S., Law, I., Page-Sharp, M., Rogerson, S., Siba, P., Ilett, K.F., Davis, T.M.E., 2009. Pharmacokinetic properties of sulfadoxine-pyrimethamine in pregnant women. Antimicrob. Agents Chemother. 53, 4368–4376. doi: 10.1128/AAC.00335-09

Kirk, D., Mittwoch, U., Stone, A.B., Wilkie, D., 1976. Limited ability of thymidine to relieve mitotic inhibition by pyrimethamine in human fibroblasts. Biochem. Pharmacol. 25, 681–685.

Kone, A., Vegte-Bolmer, M. van de, Siebelink-Stoter, R., Gemert, G.J. van, Dara, A., Niangaly, H., Luty, A., Doumbo, O.K., Sauerwein, R., Djimde, A.A., 2010. Sulfadoxine-pyrimethamine impairs *Plasmodium falciparum* gametocyte infectivity and *Anopheles* mosquito survival. Int. J. Parasitol. 40, 1221–1228. doi: 10.1016/j.ijpara.2010.05.004

Medscape, 2020. Artesunate dosing, indications, interactions, adverse effects, and more [WWW Document]. URL https://reference.medscape.com/drug/artesunate-342684 (accessed 4.21.20).

Miguel-Blanco, C., Lelièvre, J., Delves, M.J., Bardera, A.I., Presa, J.L., López-Barragán, M.J., Ruecker, A., Marques, S., Sinden, R.E., Herreros, E., 2015. Imaging-based high-throughput screening assay to identify new molecules with transmission-blocking potential against *Plasmodium falciparum* female gamete formation. Antimicrob. Agents Chemother. 59, 3298–3305. doi: 10.1128/AAC.04684-14

Müller, I.B., Hyde, J.E., 2010. Antimalarial drugs: modes of action and mechanisms of parasite resistance. Futur. Microbiol 5, 1857–1873.

Nunes, J.K., Woods, C., Carter, T., Raphael, T., Morin, M.J., Diallo, D., Leboulleux, D., Jain, S., Loucq, C., Kaslow, D.C., Birkett, A.J., 2014. Development of a transmission-blocking malaria vaccine: progress, challenges, and the path forward. Vaccine. doi: 10.1016/j.vaccine.2014.07.030

O’Neill, P.M., Barton, V.E., Ward, S.A., 2010. The molecular mechanism of action of artemisinin— the debate continues. Molecules 15, 1705–1721. doi: 10.3390/molecules15031705

Omar, M.S., Collins, W.E., Contacos, P.G., 1973. Gametocytocidal and sporontocidal effects of antimalarial drugs on malaria parasites. Exp. Parasitol. 34, 229–241. doi: 10.1016/0014-4894(73)90082-9

Paaijmans, K., Fernàndez-Busquets, X., 2014. Antimalarial drug delivery to the mosquito: an option worth exploring? Futur. Microbiol 9, 579–582. doi: 10.2217/FMB.14.30

Paton, D.G., Childs, L.M., Itoe, M.A., Holmdahl, I.E., Buckee, C.O., Catteruccia, F., 2019. Exposing *Anopheles* mosquitoes to antimalarials blocks *Plasmodium* parasite transmission. Nature. doi: 10.1038/s41586-019-0973-1

Peatey, C.L., Leroy, D., Gardiner, D.L., Trenholme, K.R., 2012. Anti-malarial drugs: how effective are they against *Plasmodium falciparum* gametocytes? Malar. J. 11, 34. doi: 10.1186/1475-2875-11-34

Peterson, D.S., Walliker, D., Wellems, T.E., 1988. Evidence that a point mutation in dihydrofolate reductase-thymidylate synthase confers resistance to pyrimethamine in *falciparum* malaria, Proc. Nati. Acad. Sci. USA.

Pigeault, R., 2015. Ecologie évolutive des interaction Hôte / Moustique / Plasmodiuml: sources d’hétérogénéité de l’infection des vecteurs. Université de Montpellier.

Pigeault, R., Vézilier, J., Cornet, S., Zélé, F., Nicot, A., Perret, P., Gandon, S., Rivero, A., 2015. Avian malaria: a new lease of life for an old experimental model to study the evolutionary ecology of *Plasmodium*. Philos. Trans. R. Soc. B Biol. Sci. 370, 20140300. doi: 10.1098/rstb.2014.0300

Plouffe, D.M., Wree, M., Du, A.Y., Meister, S., Li, F., Patra, K., Lubar, A., Okitsu, S.L., Flannery, E.L., Kato, N., Tanaseichuk, O., Comer, E., Zhou, B., Kuhen, K., Zhou, Y., Leroy, D., Schreiber, S.L., Scherer, C.A., Vinetz, J., Winzeler, E.A., 2016. High-throughput assay and discovery of small molecules that interrupt malaria transmission. Cell Host Microbe 19, 114–126. doi: 10.1016/j.chom.2015.12.001

Ramakrishnan, S.P., Basu, P.C., Harwant, S., Wattal, B.L., 1963. A study on the joint action of diamino-diphenyl-sulphone (DDS) and pyrimethamine in the sporogony cycle of *Plasmodium gallinaceum* : potentiation of the sporontocidal activity of pyrimethamine by DDS. Indian J. Malariol. 17, 141–148.

Rivero, A., Gandon, S., 2018. Evolutionary Ecology of Avian Malaria: Past to Present. Trends Parasitol. 34. doi: 10.1016/j.pt.2018.06.002

Robert, V., Awono-Ambene, H.P., Le Hesran, J.-Y., Trape, J.-F., 2000. Gametocytemia and infectivity to mosquitoes of patients with uncomplicated *Plasmodium falciparum* malaria attacks treated with chloroquine or sulfadoxine plus pyrimethamine. Am. J. Trop. Med. Hyg. 62, 210–216. doi: 10.4269/ajtmh.2000.62.210

Ruecker, A., Mathias, D.K., Straschil, U., Churcher, T.S., Dinglasan, R.R., Leroy, D., Sinden, R.E., Delves, M.J., 2014. A male and female gametocyte functional viability assay to identify biologically relevant malaria transmission-blocking drugs. Antimicrob. Agents Chemother. 58, 7292–7304. doi: 10.1128/AAC.03666-14

Sanders, N.G., Sullivan, D.J., Mlambo, G., Dimopoulos, G., Tripathi, A.K., 2014. Gametocytocidal screen identifies novel chemical classes with *Plasmodium falciparum* transmission blocking activity. PLoS One 9, e105817. doi: 10.1371/journal.pone.0105817

Saraiva, R.G., Kang, S., Simões, M.L., Angleró-Rodríguez, Y.I., Dimopoulos, G., 2016. Mosquito gut antiparasitic and antiviral immunity. Dev. Comp. Immunol. 64, 53–64. doi: 10.1016/j.dci.2016.01.015

Shinondo, C.J., Norbert Lanners, H., Lowrie, R.C.J., Wiser, M.F., 1994. Effect of pyrimethamine resistance on sporogony in a *Plasmodium berghei* /*Anophele stephensi* model. Exp. Parasitol. 78, 194–202.

Shute, P.G., Maryion, M., 1954. The effect of pyrimethamine (Daraprim) on the gametocytes and oocysts of *Plasmodium falciparum* and *Plasmodium vivax*. Trans. R. Soc. Trop. Med. an Hyg. 48, 50–63.

Sinden, R.E., 2017. Developing transmission-blocking strategies for malaria control. PLoS Pathog. doi: 10.1371/journal.ppat.1006336

Sinden, R.E., 2010. A biologist’s perspective on malaria vaccine development. Hum. Vaccin. doi: 10.4161/hv.6.1.9604

Sinden, R.E., Carter, R., Drakeley, C., Leroy, D., 2012. The biology of sexual development of *Plasmodium* : the design and implementation of transmission-blocking strategies. Malar. J. 11, 1–11. doi: 10.1186/1475-2875-11-70

Smith, D.L., McKenzie, F.E., 2004. Statics and dynamics of malaria infection in *Anopheles* mosquitoes. Malar. J. 3. doi: 10.1186/1475-2875-3-13

Sowunmi, A., Akano, K., Ntadom, G., Ayede, A., Oguche, S., Agomo, C., Okafor, H., Watila, I., Meremikwu, M., Ogala, W., Agomo, P., Adowoye, E., Fatunmbi, B., Aderoyeje, T., Happi, C., Gbotosho, G., Folarin, O., 2017. Anaemia following artemisinin-based combination treatments of uncomplicated *Plasmodium falciparum* malaria in children: temporal patterns of haematocrit and the use of uncomplicated hyperparasitaemia as a model for evaluating late-appearing anaemia. Chemotherapy 62, 231–238. doi: 10.1159/000449366

Teklehaimanot, A., Nguyen-Dinh, P., Collins, W.E., Barber, A.M., Campbell, C.C., 1985. Evaluation of sporontocidal compounds using *Plasmodium falciparum* gametocytes produced in vitro. Am. J. Trop. Med. Hyg. 34, 429–434. doi: 10.4269/ajtmh.1985.34.429

Terzakis, J.A., 1971. *Plasmodium gallinaceum* : drug-induced ultrastructural changes in oocysts. Exp. Parasitol. 30, 260–266.

Terzian, L.A., 1970. A note on the effects of antimalarial drugs on the sporogonous cycle of *Plasmodium cynomolgi* in *Anopheles stephensi*. Parasitology 61, 191–194.

Terzian, L.A., Stahler, N., Daw, A.T., 1968. The sporogonous cycle of *Plasmodium vivax* in *Anopheles* mosquitoes as a system for evaluating the prophylactic and curative capabilities of potential antimalarial compounds. Exp. Parasitol. 23, 56–66. doi: 10.1016/0014-4894(68)90042-8

Van Eijk, A.M., Hill, J., Alegana, V.A., Kirui, V., Gething, P.W., ter Kuile, F.O., Snow, R.W., 2011. Coverage of malaria protection in pregnant women in sub-Saharan Africa: a synthesis and analysis of national survey data. Lancet Infect. Dis. 11, 190–207. doi: 10.1016/S1473-3099(10)70295-4

Vézilier, J., Nicot, A., Gandon, S., Rivero, A., 2015. *Plasmodium* infection brings forward mosquito oviposition. Biol. Lett. 11, 2–6. doi: 10.1098/rsbl.2014.0840

Vézilier, J., Nicot, A., Gandon, S., Rivero, A., 2012. *Plasmodium* infection decreases fecundity and increases survival of mosquitoes. Proc. Biol. Sci. 279, 4033–41. doi: 10.1098/rspb.2012.1394

Vézilier, J., Nicot, A., Gandon, S., Rivero, A., 2010. Insecticide resistance and malaria transmission: infection rate and oocyst burden in *Culex pipiens* mosquitoes infected with *Plasmodium relictum*. Malar. J. 9. doi: 10.1186/1475-2875-9-379

Vincke, I.H., 1970. The effects of Pyrimethamine and Sulphormethoxine on the pre-erythrocytic and sporogonous cycle of *Plasmodium berghei berghei*. Ann. Soc. Belg. Med. Trop. (1920). 50, 339–358.

Wadi, I., Anvikar, A.R., Nath, M., Pillai, C.R., Sinha, A., Valecha, N., 2018. Critical examination of approaches exploited to assess the effectiveness of transmission-blocking drugs for malaria. Future Med. Chem. 10, 2619–2639. doi: 10.4155/fmc-2018-0169

Wadi, I., Nath, M., Anvikar, A.R., Singh, P., Sinha, A., 2019. Recent advances in transmission-blocking drugs for malaria elimination. Future Med. Chem. 11, 3047–3088. doi: 10.4155/fmc-2019-0225

WHO, 2019. World Malaria Report 2019. Geneva.

Zélé, F., Nicot, A., Berthomieu, A., Weill, M., Duron, O., Rivero, A., 2014. *Wolbachia* increases susceptibility to *Plasmodium* infection in a natural system. Proc. R. Soc. B 281. doi: 10.1098/rspb.2013.2837

Zhou, G., Kohlhepp, P., Geiser, D., Frasquillo, M. del C., Vazquez-Moreno, L., Winzerling, J.J., 2007. Fate of blood meal iron in mosquitoes. J. Insect Physiol. 53, 1169–1178. doi: 10.1016/j.jinsphys.2007.06.009

